# Wild birds’ plight and role in the current bird flu panzootic

**DOI:** 10.1101/2023.05.02.539182

**Authors:** Marcel Klaassen, Michelle Wille

## Abstract

The current avian influenza panzootic is unprecedented and catastrophic for birds. With a focus on the implications of this panzootic for poultry, there is limited attention on wild birds. We highlight shortcomings and geographic biases in reporting leading to a severe underappreciation of wildlife mortality. We estimate the scale of mortality amongst wild birds is in the millions rather than tens-of-thousands reported, through comparison of notification data to accounts literature. The outbreaks amongst wild birds are causing population and species level concerns which may drive extinctions and jeopardise decades of conservation efforts.

## Main

Since its emergence in Hong Kong in 1996, high pathogenicity avian influenza (HPAI) viruses of the A/Goose/Guangdong/1/96 lineage have steadily evolved. Since 2014, they have become increasingly widespread and progressively more destructive across the poultry industry and wild birds, globally ^1^. This has culminated in the emergence of the descendant lineage 2.3.4.4b, which since October 2021 has spread to all continents except the Antarctic and Australia, resulting in a panzootic of unprecedented magnitude (Fig 1A-B). According to the World Animal Health Information System (WAHIS) from the World Organisation for Animal Health ^2^ this panzootic has resulted in the death and destruction of more than half a billion poultry. Wildlife, notably birds, have also taken a profound hit. Based on available data, 2.3.4.4b has infected and killed a substantially greater diversity of wild birds (i.e., 320 species belonging to 21 orders) compared to previous lineages (Fig. 1 C). Moreover, while prior to October 2021 most HPAI cases in wild birds occurred near intensive poultry production, recent outbreaks in wild birds now also include remote areas thousands of kilometres away from poultry production.

**Figure 1.**
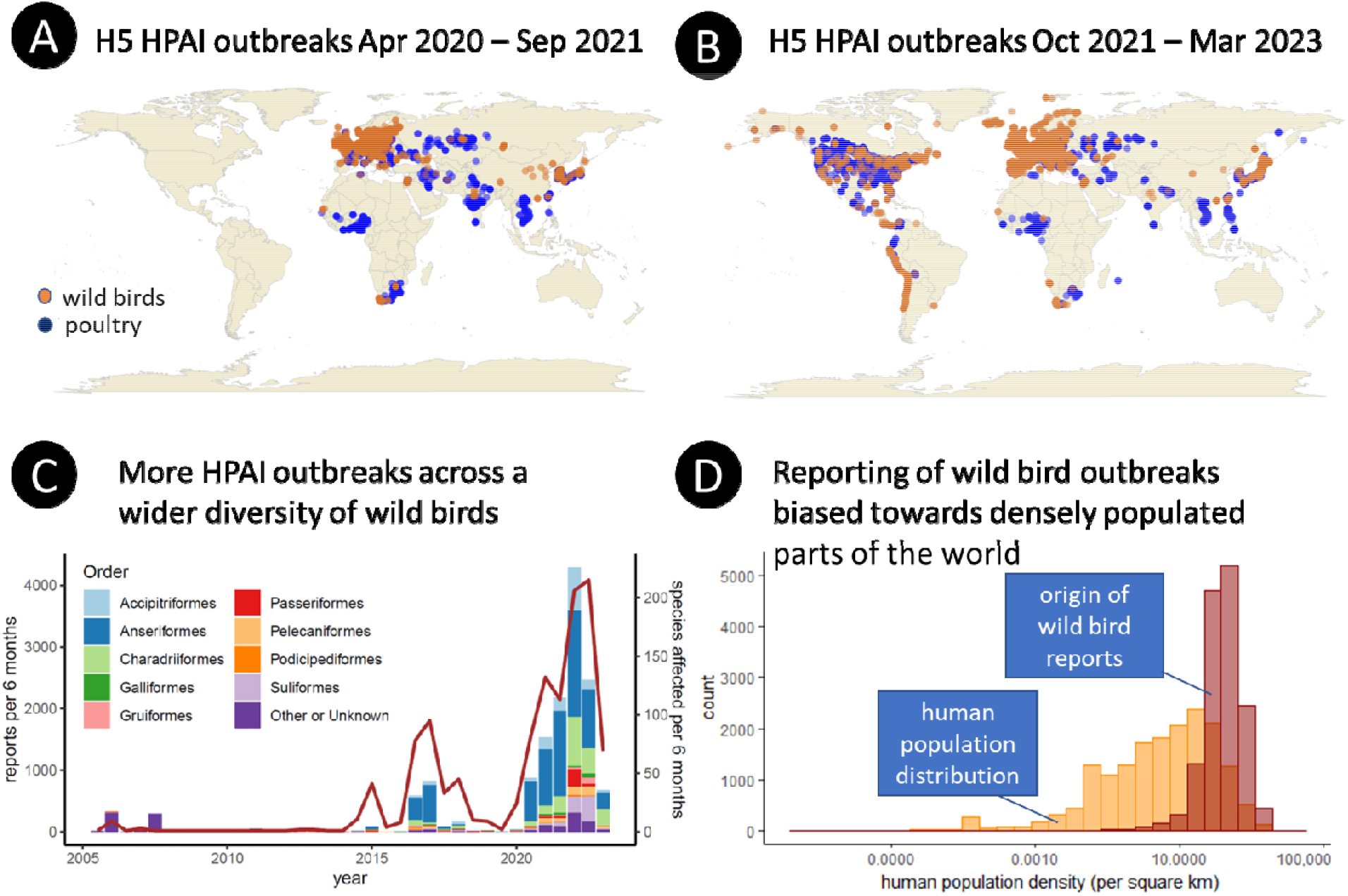
(**A**) HPAI H5 outbreaks in wild birds and poultry in the two waves prior to the current panzootic. (**B**) HPAI H5 outbreaks during the current panzootic. (**C)** Number of reports of HPAI in wild birds, per half year, across avian orders (stacked bars, left axis), and the number of wild bird species involved (brown line, right axis). (**D)** Histogram of the number of reports received on wild birds infected with HPAI since 2005 as a function of human population density (on a 10-log scale) compared to a histogram depicting the global distribution of human density across the land masses (using a one-by-one degrees of latitude and longitude gridded map) (orange).

While the reported scale of poultry mortality due to HPAI is likely aligned to reality, the 68,013 wild bird casualties and 289 mammalian cases reported in WAHIS^2^ are not. In part, this is due to inconsistent reporting, either including all observed carcasses, only those tested and confirmed, or no information on numbers at all. For instance, the outbreaks comprising 24,463 Cape Cormorants in South Africa (*Phalacrocorax capensis*) and 9,029 Common Cranes (*Grus grus*) in Israel appear to be reported in full. Apparently more widespread is the reporting of fractions of cases compared to what is stated in the media ^3^, or from publications. For example, in the WAHIS database there are 41 HPAI notifications for Sandwich Terns (*Thalasseus sandvicensis*) involving 68 individuals globally, since 1 October 2021. This contrasts sharply with 9,600 dead Sandwich Terns ^4^ reported in the Netherlands alone, of which only 19 are included in WAHIS. In the media a further 7000 Sandwich Tern carcasses were reported in France, the UK and Germany, which together comprises a two to three order of magnitude difference between official notifications in WAHIS and reports from the ground. Similarly, 73 dead Great Skuas (*Stercorarius skua*) have thus far been reported to WAHIS, while 1400 dead individuals were found on the small Scottish island of Foula alone ^5^. Additionally, 593 Dalmatian Pelicans (*Pelecanus crispus*) were reported to WAHIS from Greece, whereas 2,286, or approximately 40% of the SE European population, have been reported in the literature ^6^. Finally, in some cases, outbreaks are reported only in the literature and not in WAHIS, such as the 2140 Great White Pelicans (*Pelecanus onocrotalus*) found dead in Mauritania in early 2021 ^7^.

Humans report outbreaks, and the correlation between the distribution of wildlife cases reported and human population density strongly suggests the potential for vast underreporting in areas where few people live (Figure 1D). Combined with the high underreporting of outbreaks in areas where human population densities are still high (e.g., the case of the Dutch Sandwich Terns), suggests only a fraction of outbreaks in wildlife have been detected and appropriately reported. Therewith, the number of wild birds impacted is conceivably in the millions rather than the tens of thousands that have been reported and collated in WAHIS ^2^.

If HPAI continues to frequently spill over from poultry to wildlife, wild bird populations will continue to be affected. The global poultry population currently comprises 70% of the world’s avian biomass ^8^ and plays a central role in the perpetuation of HPAIs ^9^. Until recently, HPAI outbreaks in wild birds could be directly linked to spill-over from poultry, and wild birds did not play a central role in virus perpetuation. For example, HPAI repeatedly disappeared after annual incursions in Europe between 2005 and 2008, and North America and Europe in 2014 ^10^. However, now HPAI 2.3.4.4b has adapted from poultry to wild birds ^11^, explaining the explosion of sustained outbreaks in wild birds since 2020, including outbreaks in remote areas with no poultry.

Despite not historically being the main reservoir for HPAI, wild birds play a key role in long distance viral spread. This is facilitated by individuals that show only mild or no clinical signs after virus infection. Tracking studies in China have shown that Mallards (*Anas platyrhynchos*) are capable of flying hundreds of kilometres while infected with HPAI 2.3.4.4b ^12^. These birds’ tolerance for HPAI infection is likely explained by their infection history. Waterfowl typically have high rates of low pathogenic avian influenza prevalence, including exposure to low pathogenic H5 strains ^13^. These infections may provide homo- or heterosubtypic immunity against subsequent HPAI infections ^14^, preventing infection or dampened disease severity ^14^, serving as one explanation how such virulent virus can spread so readily on-board wild birds.

Given the enormous burden of this virus on poultry and wild birds alike, there are increased calls for poultry vaccination. While proven successful in significantly reducing mortality ^15^, vaccination of poultry may also drive virus evolution ^16^, leading to continued virus circulation in vaccinated flocks with limited disease signs^17^, contributing to endemicity and spill over risk to wild birds. However, while not a silver bullet, when done in combination with monitoring to guarantee disease freedom, a “vaccination plus strategy” should ideally decrease not only disease burden on poultry, but also on wild birds and mammals by limiting spill-over from poultry production.

In summary, despite lacking an accurate estimate of the true impact on wildlife, we are witnessing a panzootic of an unprecedented and enormous scale. This panzootic did not emerge from nowhere, but rather is the result of 20 years of viral evolution in the ever-expanding global poultry population. Given the key role of poultry production in food chains, and the effect on livelihoods, it is logical for countries to prioritize their response towards poultry. However, that wild bird outbreaks are widely neglected, to the degree that we do not even know the order of magnitude of deaths, nor the population and ecosystem consequences, is highly concerning. As a result, the true impact of this panzootic on wild birds may not be recognised for years to come, and some species may never recover.

## Online Methods

Data were downloaded from the World Organization for Animal Health (WOAH) World Animal Health Information System (WAHIS) database on 05/03/2023. Data extracted by Marcel Klaassen, Deakin University. Reproduced with permission. WOAH bears no responsibility for the integrity or accuracy of the data contained herein, but not limited to, any deletion, manipulation, or reformatting of data that may have occurred beyond its control.

Special thanks to Paolo Tizzani from WOAH for kindly providing additional information on this data source.

“Al indeterminatum fau” are assumed to be birds.

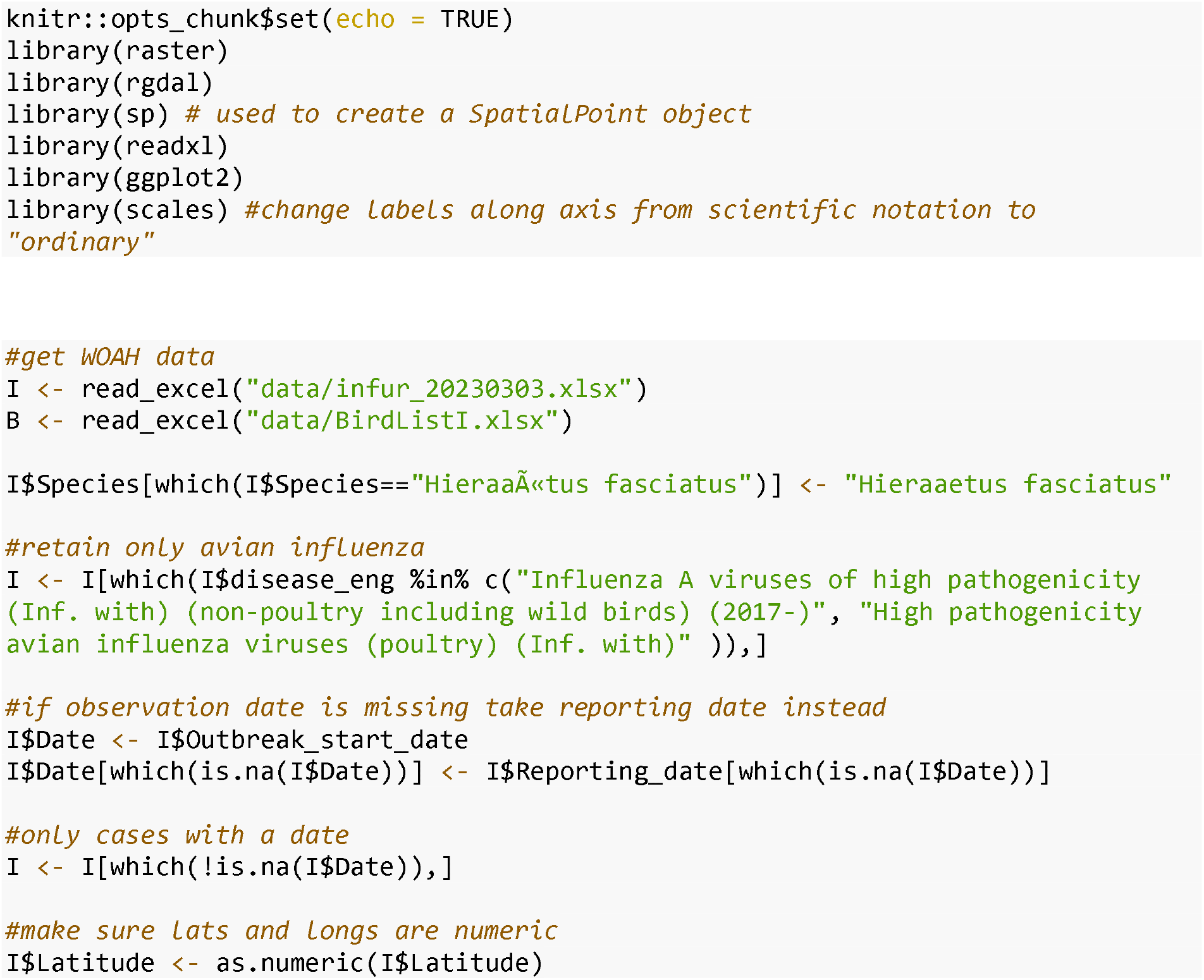
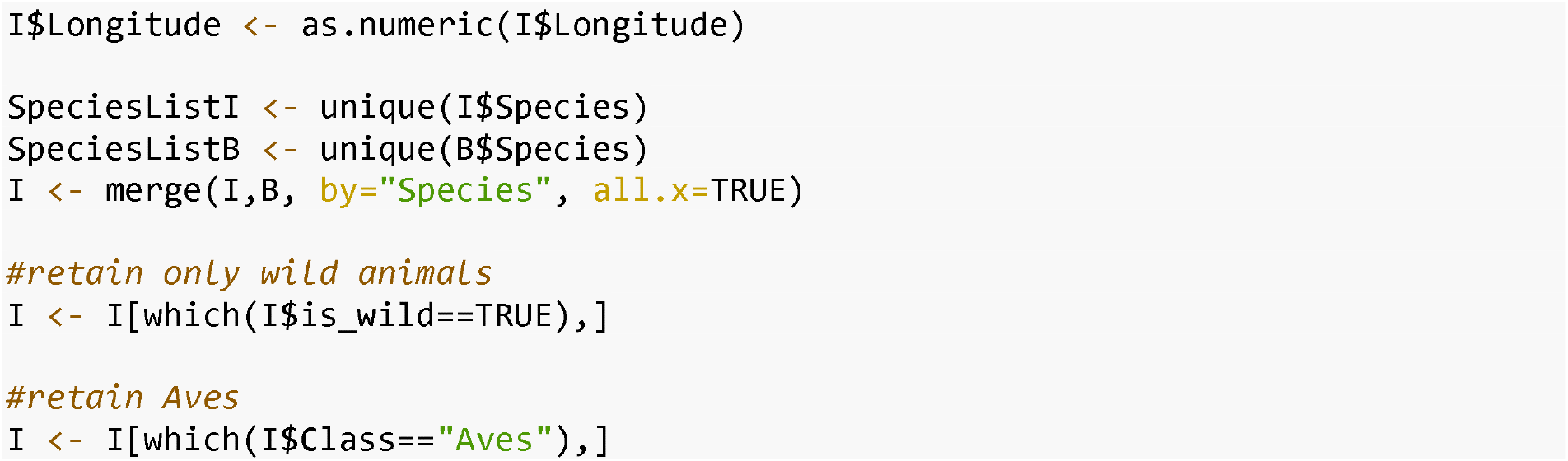

Finding wild birds affected by HPAI may be a function of the number of people living there. Large colonies of (sea) birds can often be found in remote areas. The question thus arises what we actually do pick up of what is happening to wild birds globally?

World population density data was downloaded from http://sedac.ciesin.columbia.edu/data/collection/gpw-v4/documentation and plotted un-transformed and on a log scale. From these maps and for all HPAI incidence data we extracted the corresponding human population density of which next a histogram was plotted.This histogram suggests that the majority of reports come from areas where human population densities are at an intermediate level. This pattern may be due to the fact that where many people live few birds occur in high densities and where few people live there may be affected birds, but they are less likely to be noted and reported.

Further support for this pattern emerges when studying the outbreak maps and focusing on the northern hemisphere shorelines of temperate and Arctic regions. These are characterised by large seabird colonies in which large outbreaks have been recorded over the past year, but only in those areas where population densities are relatively high (Svalbard, Iceland, Europe, west coast of Canada and south coast Alaska) but not elsewhere (north coast North America, Greenland, Russia). It should also be noted that due to the COVID-19 pandemic research activities in these regions have also been limited.

There is thus a very high chance that the size of the AIV panzootic is considerably under-reported.

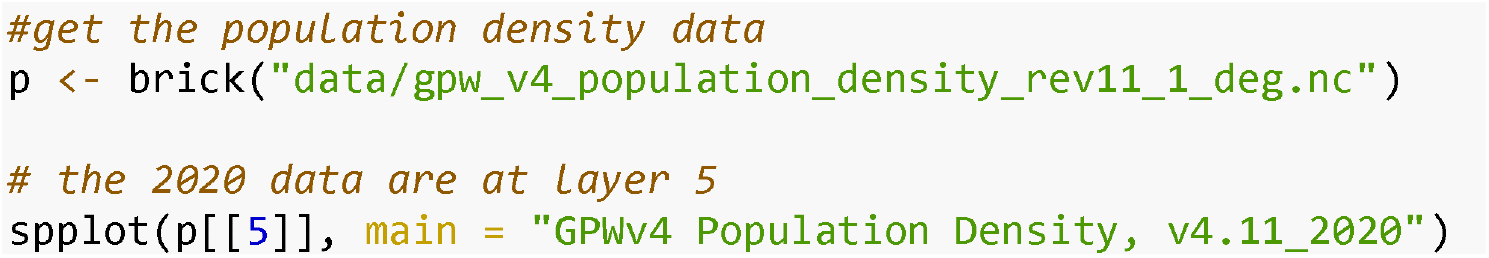

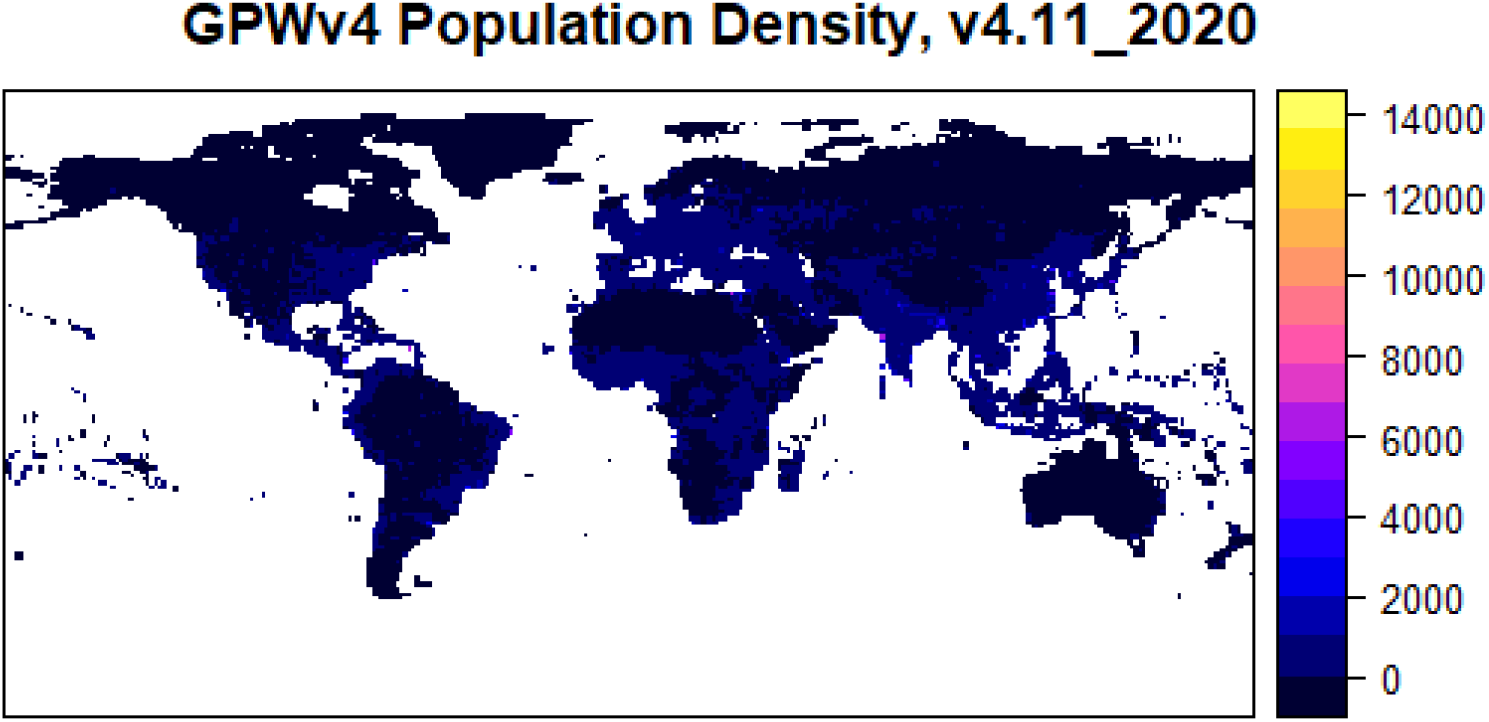

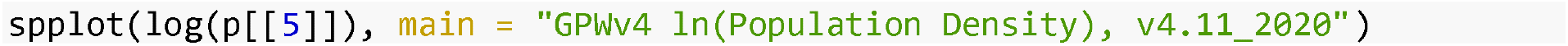

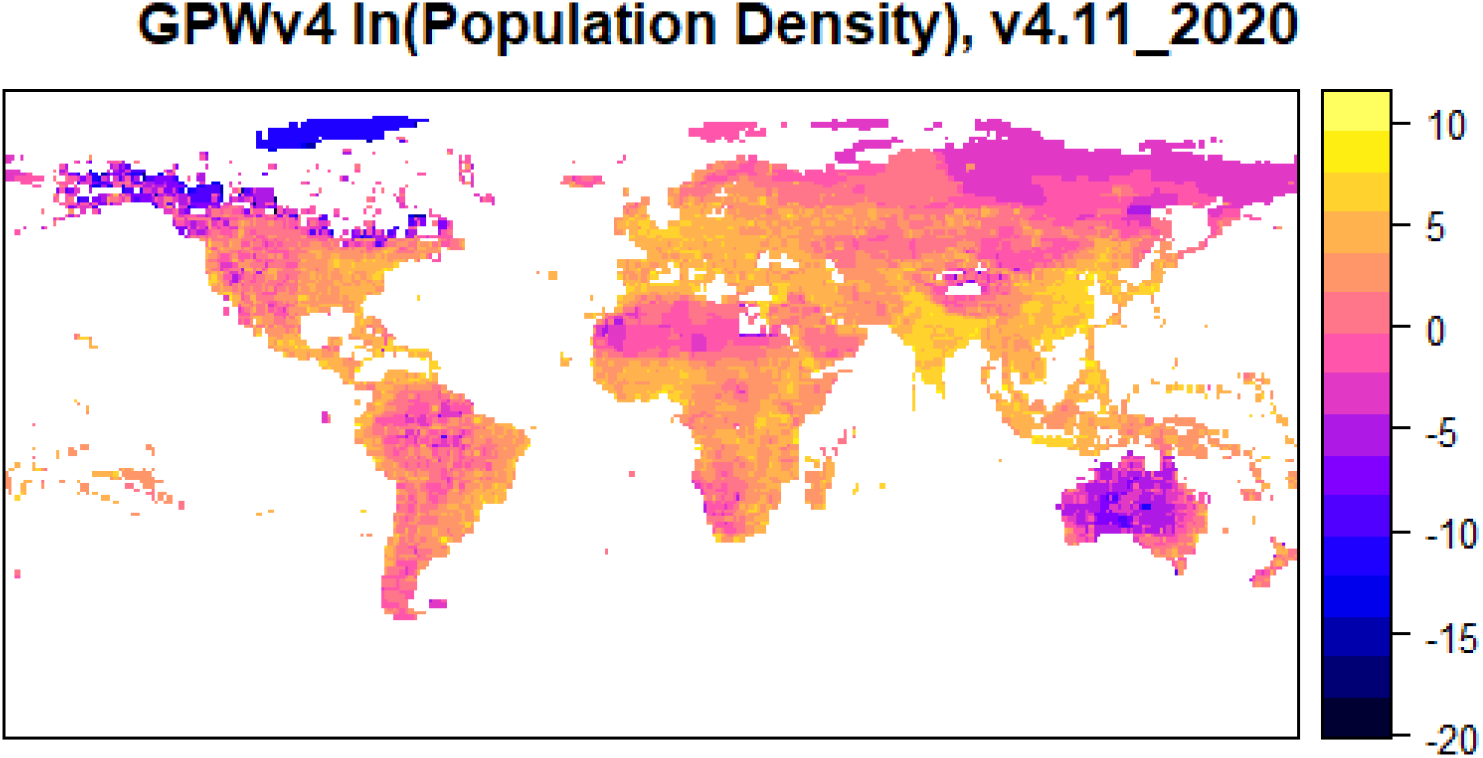

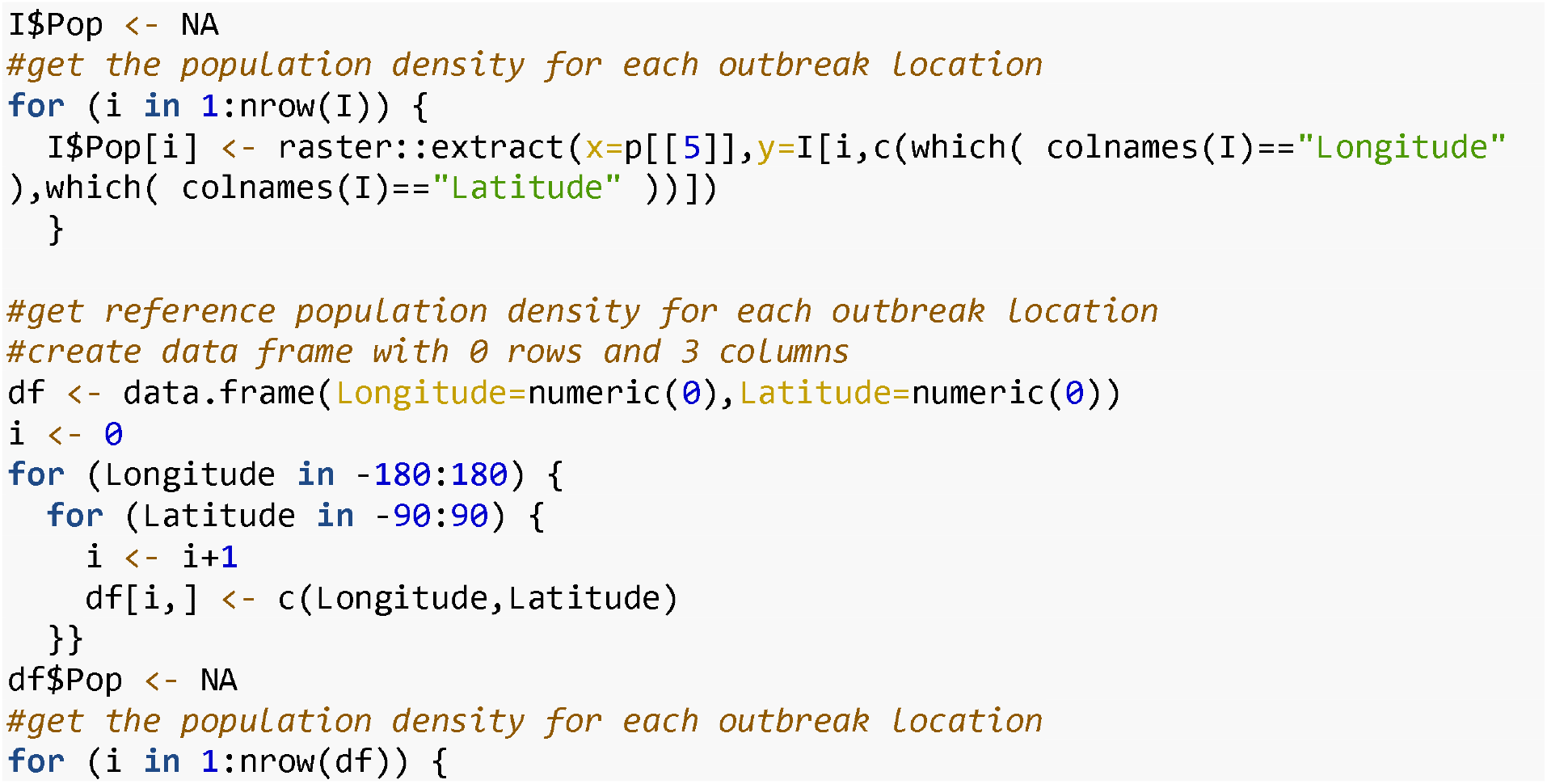
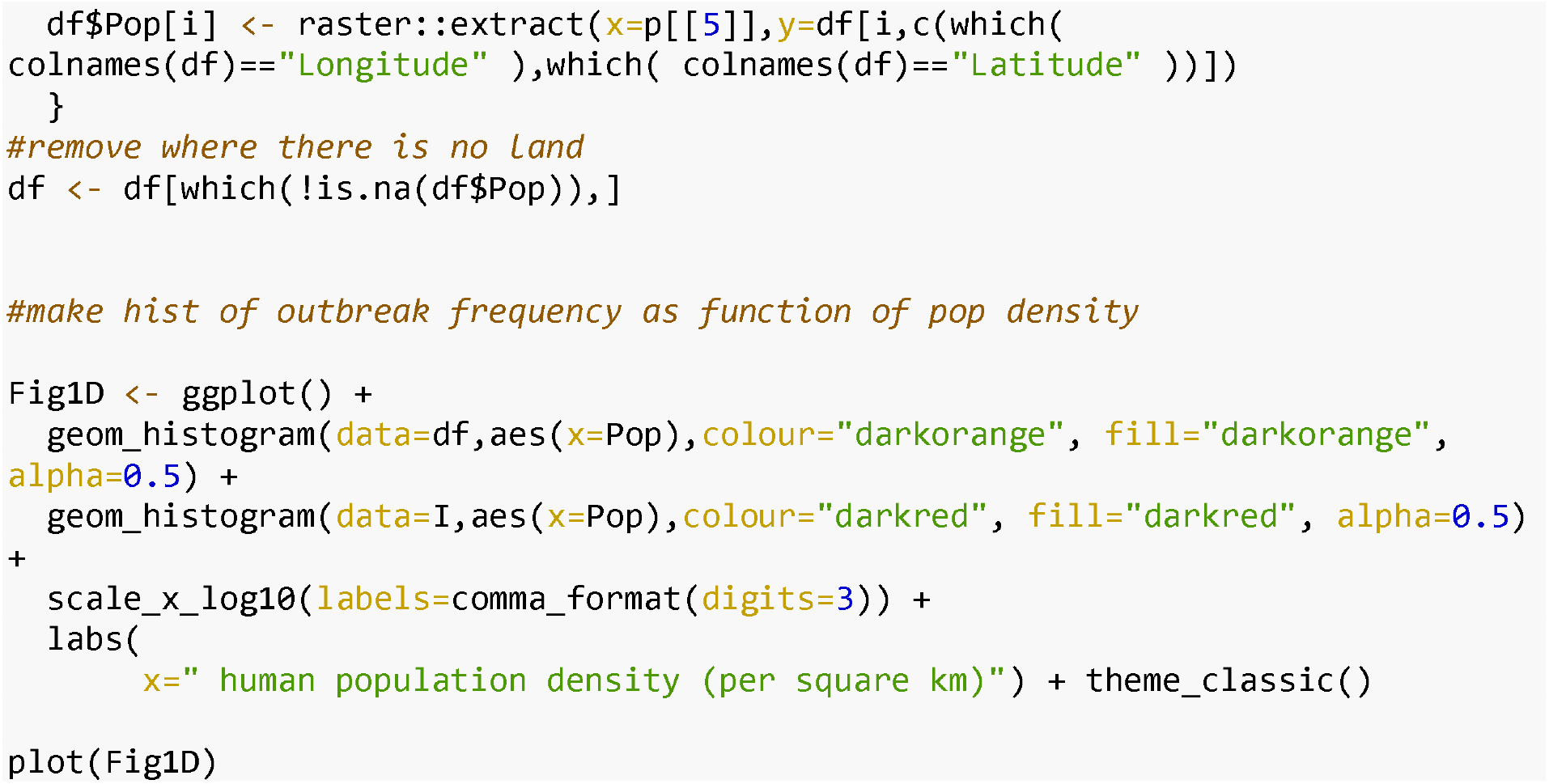

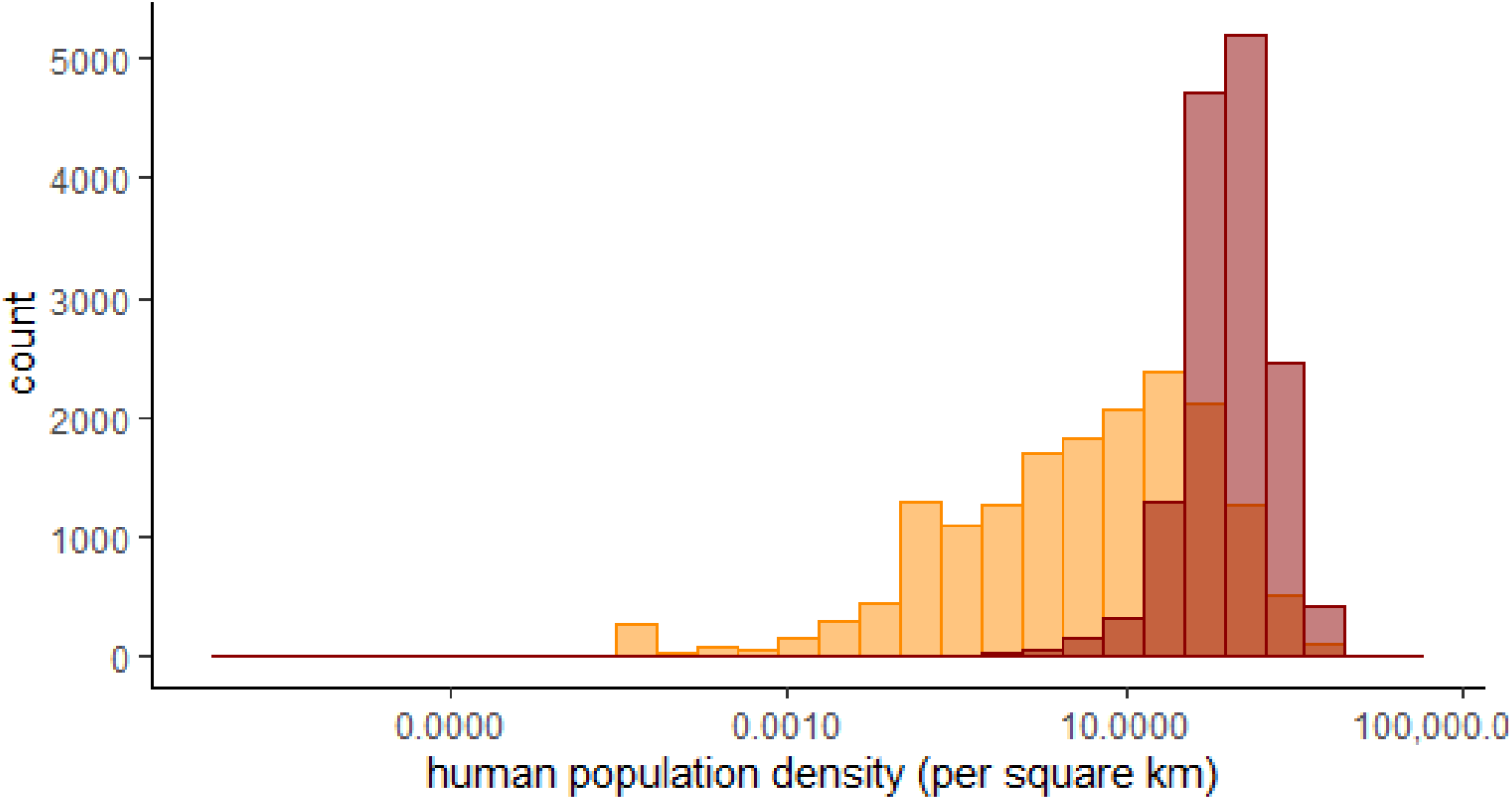

## Acknowledgements

Research in Marcel Klaassen’s lab is undertaken with support from Australian Research Council (ARC) Discovery Project Grant DP19010186. Michelle Wille is funded by an ARC Discovery Early Career Research Award (DE200100977). The WHO Collaborating Centre for Reference and Research on Influenza is funded by the Australian Department of Health.

